# Effect of dopamine D2 blockade on behavioural and electrophysiological measures of inhibitory control

**DOI:** 10.64898/2026.09.16.752057

**Authors:** V. Rohira, J.P. Coxon, K. Stevens, R. Joshi, N. Thirugnanasambandam, I. Leunissen, T.T.-J Chong, M.A. Bellgrove

**Affiliations:** The Turner Institute for Brain and Mental Health, School of Psychological Sciences, Monash University, Victoria 3800, Australia; Department of Biosciences and Bioengineering, Indian Institute of Technology Bombay, Powai, Mumbai 400076; Department of Cognitive Neuroscience, Faculty of Psychology and Neuroscience, Maastricht University, Maastricht 6229 EV, The Netherlands; Maastricht Brain Imaging Centre (MBIC), Maastricht University, Oxfordlaan 55, Maastricht 6229 EV, The Netherlands; Department of Neurology, Alfred Health, Melbourne, Victoria 3004, Australia; Department of Clinical Neurosciences, St Vincent’s Hospital, Melbourne, Victoria 3065, Australia

**Keywords:** Response inhibition, Proactive inhibition, Reactive inhibition, Dopamine, EEG, Stop-signal task

## Abstract

Inhibitory control is an essential executive function that encompasses both the stopping of an action in response to an unexpected stop signal (reactive response inhibition) and slowing of responses in anticipation of stopping (proactive response inhibition). Fronto-basal-ganglia circuits are thought to differentially contribute to response inhibition, with the indirect pathway implicated in proactive inhibition. Given the high expression of dopamine D2 receptors in the indirect pathway, pharmacological blockade of D2 receptors is expected to impact proactive inhibition, as indicated by response times on Go trials. The current study aimed to investigate this experimentally and study the effect of 600 mg of sulpiride, a D2 receptor antagonist, on behavioural and electrophysiological measures of proactive and reactive inhibition in humans. Human participants (N=24) completed an anticipated response version of the stop-signal task on either sulpiride or placebo in a double-blind within-subjects design. Sulpiride led to increased variability of Go response times and attenuation of the frontocentral negativity/readiness potential, considered to be a signature of proactive inhibition. In contrast, sulpiride had no effect on stop-signal reaction time or event-related potential components associated with reactive inhibition. These findings suggest that D2 receptor blockade selectively alters processes involved in proactive inhibition while leaving reactive inhibition unaffected, consistent with a key role for the indirect basal ganglia pathway in proactive response inhibition.

## Introduction

Response inhibition is a core executive function and is essential for flexible goal-directed behaviours in everyday life. A distinction is made between *reactive inhibition* and *proactive inhibition*. Reactive inhibition refers to the ability to rapidly stop an already initiated action following the presentation of an unexpected stop signal, whereas proactive inhibition involves a preparatory process that may facilitate movement cancellation in anticipation of a stop signal [Aron, 2011] by altering the dynamics of movement preparation. Response inhibition is commonly studied using stop-signal tasks with reactive inhibition performance quantified via the stop-signal reaction time (SSRT) [Aron, 2007; Verbruggen et al., 2019]. In contrast, proactive inhibition is inferred from a subtle slowing of Go trial responses in conditions with higher stop-signal probability, likely the result of increasing decision uncertainty and dynamic competition between the direct and indirect basal ganglia pathways [Dunovan and Verstynen, 2016]. Notably, neuroimaging research implicates fronto-basal-ganglia pathways in both reactive and proactive inhibition [Aron et al., 2014; Coxon et al., 2016; Dunovan et al., 2015; Leunissen et al., 2016; Zandbelt and Vink, 2010].

Cortical input to the basal ganglia can occur via distinct functional pathways; namely ‘direct’ and ‘indirect’ projections involving the striatal D1 and D2 receptors respectively, and a ‘hyperdirect’ projection from cortex to the subthalamic nucleus (STN). These pathways exert opposing facilitatory (direct pathway) and inhibitory (indirect, hyperdirect pathway) effects on the thalamocortical projection and are implicated in the selection, initiation, and inhibition of actions [Jahanshahi et al., 2015; Schmidt et al., 2013]. The hyperdirect pathway is thought to be critical for reactive inhibition, whereas the indirect pathway is implicated in proactive inhibition [Aron, 2011], although such distinctions in humans remain controversial [Wessel, 2026]. Pharmacological challenge via drugs acting on specific receptors (e.g. dopamine D1 or D2 receptors) might allow for specific inferences regarding the contribution of cortico-basal ganglia pathways to response inhibition.

Multiple studies in clinical and non-clinical populations have shown that indirect catecholamine (dopamine and noradrenaline) agonists, such as methylphenidate and atomoxetine, improve reactive stopping [Chamberlain et al., 2006; Nandam et al., 2011a], although less is known about receptor level effects. Nandam et al. [2013] observed enhanced reactive stopping in healthy individuals on cabergoline, a high affinity D2 receptor agonist, in comparison to placebo. Moderate deficits in reactive stopping have been reported in Parkinson’s disease patients off medication, which are improved with dopaminergic medication in the earlier stages of the disease [Manza et al., 2017]. Evidence from positron emission tomography (PET) studies provides further support for a role of D2/D3 receptors in reactive stopping. D2/D3 receptor availability in the dorsal striatum has been reported to be negatively correlated with SSRT, but not with go-trial reaction time, suggesting that D2/D3 receptor availability may be specifically associated with the ability to inhibit an ongoing motor response instead of a more general enhancement of motor performance [Ghahremani et al., 2012; Robertson et al., 2015]. The above evidence suggests a potential role for dopamine in response inhibition processes in humans, but the exact role of dopamine D2 receptors remains unanswered.

Pharmacological studies of proactive inhibition are limited. Ropinirole is a D2/D3 receptor agonist that binds with high affinity at the dopamine D3 receptor [Perachon et al., 1999]. Ropinirole was shown to reduce proactive response inhibition without any changes in reactive stopping [Rawji et al., 2020]. Another study has highlighted that ropinirole’s effect on reactive stopping depends on an individual’s dopamine neurotransmission genetic profile. Ropinirole was shown to improve reactive inhibition compared to placebo in healthy adults with genetic DNA variations favouring lower basal dopamine neurotransmission but impair reactive inhibition in adults with DNA variations favouring higher basal dopamine neurotransmission [MacDonald et al., 2016]. The above evidence suggests a role for dopamine in response inhibition processes in humans.

Work in rodents also suggests a role for dopamine receptors in reactive stopping. Sulpiride is a dopamine D2 receptor antagonist [Caley and Weber, 1995] and has been shown to increase both SSRT and go-trial reaction time in rats, whereas a dopamine D1 receptor antagonist decreased SSRT without any changes in proactive inhibition. [Eagle et al., 2011]. These effects were evidenced specifically when dopamine D1 and D2 receptor antagonists were administered through infusion in the dorsomedial striatum, but not the nucleus accumbens core, or via intraperitoneal injections [Bari and Robbins, 2013]. However, findings from rodent models may not translate to primates due to task factors and whether comparable effects of D2 receptor blockade on proactive and/or reactive inhibitory control in humans are unknown.

Human electroencephalography (EEG) studies have identified event-related potentials (ERP) associated with response inhibition. The Contingent Negative Variation (CNV) is a slow negative ERP that develops during the anticipatory interval between a warning stimulus and a subsequent imperative stimulus requiring a motor response [Walter et al., 1964]. The late component of the CNV reflects processes related to attentional and motor preparation [Rohrbaugh et al., 1976; Tecce, 1972]. Importantly, studies have reported larger late CNV amplitude in Go trials with a higher probability of an upcoming stop signal as compared to Go trials with a lower stop-signal probability [Brevers et al., 2020; Lee and Kang, 2020]. Consequently, the CNV has been proposed as a biomarker of proactive control during response inhibition [Brevers et al., 2020].

EEG signatures of response inhibition following a stop-signal have also been studied and include the N2 and P3 peaks of the ERP [De Jong et al., 1990; Huster et al, 2013; Waller et al., 2019; Zhu et al., 2024]. Additionally, a prominent P1 has been observed in some participants [Wessel and Aron, 2015]. Despite the emphasis on P3 in the literature, its functional significance remains under debate. Kok et al. [2004] proposed that successful response inhibition may rely on the N2/P3 being early enough following a stop-signal to cause rapid downstream effects on activation processes. This idea has been supported by studies that show earlier P3 onset latency on successful stop trials as compared to unsuccessful stop trials [Diesburg and Wessel, 2015; Dykstra et al., 2020; Hervault et al., 2025; Wessel and Aron, 2015]. However, for the P3 to be causally involved in reactive inhibition, the onset latency should presumably precede the stopping latency, which is typically not what is observed [Huster et al., 2020; Thunberg et al., 2020]. Moreover, while P3 latencies have been associated with SSRT, this association is not specific solely to the P3. N2 latencies precede the P3 by about 150 ms and have also been associated with SSRT [Huster et al., 2020]. It therefore remains unclear whether the P3 is causally involved in implementing reactive inhibition or is related more strongly to processing awareness of success or error on a given trial.

The present study aimed to investigate the effects of dopamine D2 receptor blockade, using sulpiride, on behavioural and EEG signatures of proactive and reactive inhibition in humans. Previous studies indicate that the effect of sulpiride on dopaminergic transmission is dose-dependent. At lower doses (e.g., 200mg), sulpiride influences presynaptic D2 autoreceptors and thereby increases dopamine release [Jocham et al., 2011], whereas higher doses result in greater post-synaptic dopamine D2 receptor blockade [Eisenegger et al., 2014]. We administered a 600 mg dose of sulpiride, which antagonises the post-synaptic dopamine D2 receptor and impairs dopamine transmission [Mehta et al., 2007; Takano et al., 2006].

In this study, a stop-signal paradigm employing an anticipated response for the Go task was utilised [Coxon et al., 2006; Leunissen et al., 2017], with a task manipulation similar to [Zandbelt and Vink, 2010]. Visual cues signalled the probability of a stop-signal occurring from one trial to the next. Based on functional neuroimaging studies showing striatal activation during proactive inhibition [Leunissen et al., 2016; Zandbelt and Vink, 2010] the study sought to understand the role of D2 receptors, and by extension the indirect cortico-basal ganglia pathway, in proactive inhibition. In accordance with the rodent study by [Eagle et al., 2011], it was hypothesised that sulpiride would impair reactive inhibition, evidenced by increased SSRTs. Further, it was hypothesised that sulpiride would increase proactive inhibition, seen as an increase in Go reaction time in trials with higher stop-signal probability. With regard to sulpiride’s effects on ERPs, it was expected that sulpiride would reduce N2 and P3 amplitudes following the stop-signal (reactive inhibition) due to impaired reactive inhibition. Further, a larger CNV amplitude on Go trials that have a higher probability of a stop-signal occurring was expected, and that sulpiride would alter fronto-central CNV amplitude.

## Methods

This study employed a randomised, double-blind, placebo-controlled, within-subjects cross-over design of the effect of sulpiride (600 mg) on the anticipated response stop-signal task. Eligible participants were enrolled in the study and randomised to either a “sulpiride first” or “placebo first” condition according to a computerised sequence generated by an external randomisation service (Griffith University). Study drugs were compounded by an external compounding pharmacy (Stenlake Compounding Pharmacy) and provided in identical blue, gelatine filled capsules of either sulpiride, 600mg, or microcrystalline cellulose (placebo) and marked only with the participant’s study and session numbers. EEG data was recorded while participants completed the anticipated response stop-signal task. Behavioural and EEG data analysis was performed blinded to Drug session using custom Matlab scripts. Unblinding was performed after all data collection and individual subject/session analyses were complete.

### Participants and drug administration protocol

Participants were excluded if they reported any history of neurological or psychiatric illness; head injury involving loss of consciousness; electroconvulsive therapy; use of psychotropic medication; significant illicit drug use (defined as greater than 5 lifetime intake of any illicit drug except cannabis; or more than monthly cannabis intake); any illicit drug use in the 6 months before testing; current or significant past history of tobacco smoking (defined as greater than 5 cigarettes per week) or alcohol dependence (defined as greater than 24 units a week). Female participants were required to be taking the combined estrogen/ progesterone oral contraceptive pill and to return a negative pregnancy test. Participants underwent a medical and psychiatric screen by a consultant neurologist (TTJC) using the M.I.N.I. Screen [Lecrubier et al., 1997] and the Kessler K10 [Kessler et al., 2003], and were excluded if these measures indicated elevated symptoms consistent with a current psychiatric disorder or if there were any contraindications to sulpiride. All participants gave written informed consent, and all procedures were approved by the ethical review board of Monash University, Melbourne. Ethical guidelines were in accordance with the Declaration of Helsinki.

Participants attended two sessions, on the same day and at the same time, one week apart. They were required to refrain from alcohol for 12 hours before testing and from any food or drink for 1 hour before attending the study session. On arrival, heart rate and blood pressure were taken and, after confirming both were in the normal range, participants were asked to ingest the study capsule with a small amount of water. Participants were not permitted to eat or drink (other than small amounts of water) until the study session ended, to avoid interference with drug absorption. Sulpiride is absorbed slowly from the gastrointestinal tract [Wishar et al., 2006] and, at doses above 400mg, takes 3 hours to reach peak plasma levels [Takano et al., 2006]. Participants waited in the testing lab or an adjacent room and were permitted to read and relax for approximately 2 hours following drug administration, after which EEG preparation and impedance checking commenced. Heart rate and blood pressure were measured and subjective ratings of alertness were assessed at baseline and each hour following drug administration, using the Bond-Lader Visual Analogue Scales (VAS) of alertness, calmness and contentedness [Bond and Lader, 1974]. A drug effects questionnaire (neurovegetative list, NVL) [Rush et al., 2003] was administered 3 hours after drug administration and again when testing was complete. The questionnaire contained 20 items (examples included ‘any effect’, ‘bad effect’, ‘performance impaired’, ‘fatigued’) and participants responded verbally on a scale of 1 to 5 (1 = not at all; 2 = a little bit; 3 = moderate; 4 = quite a bit; 5 = very much). A drug allocation awareness check was conducted at conclusion of the second study session, with participants reporting which session they believed they had ingested sulpiride and how confident they were in their ratings (out of 10) [Eisenegger et al., 2010].

### Materials and task procedures

Participants were seated in a darkened room 56 cm from a 21-inch CRT monitor (85 Hz, 1024 x 768 pixel resolution), with their chin and forehead resting in a stationary head mount, and asked to perform a variant of the anticipated response stop-signal task [Coxon et al., 2007; Leunissen et al., 2017; Zandbelt et al., 2010]. The paradigm was run using custom code written in LabVIEW (National Instruments). Participants viewed a vertical indicator with a horizontal target line situated at 4/5ths of the indicator length. Each trial began with a warning cue for 500 ms (a black rectangle surrounding the indicator in the participants peripheral vision). Subsequently the indicator began to fill with colour, at a constant rate from the bottom towards the top in 1 second (Figure 1). The primary task was to stop the indicator at the target (800 ms from onset) by releasing a mouse key via a right index finger extension movement (Go trials). Go trials were to be performed as accurately and consistently as possible.

**Figure 1.**
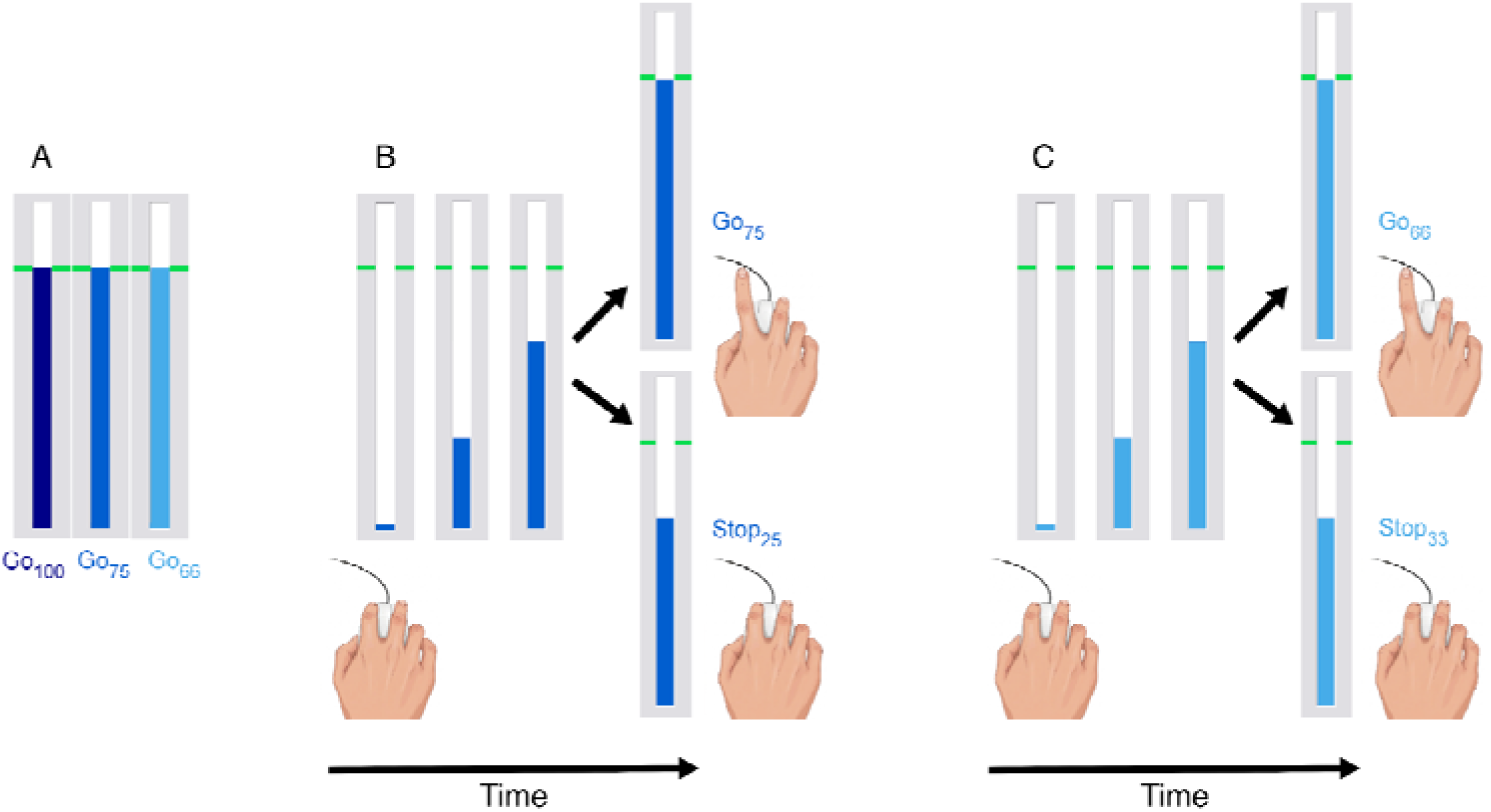
Behavioural task. (A) The indicator colour conveyed information about the likelihood the trial would be a Go/Stop trial. Dark (navy) blue, medium (royal) blue, and light blue were associated with Go trial probabilities of 100%, 75%, and 66%. (B,C) Participants viewed the indicator fill at a constant rate. Participants were required to release a mouse key to stop the indicator as close to the horizontal target line as possible (Go trial), but to inhibit this response if the filling indicator stopped automatically prior to reaching the horizontal target line (Stop trial).

To measure proactive inhibition, we manipulated the indicator fill colour from trial-to-trial (Figure 1A). The following text was fixed to the top of the CRT monitor for the participant’s reference throughout the experiment:

> “The darker the bar, the more likely it will be a Go trial.
>
> The lighter the bar, the more likely it will be a Stop trial”If the filling indicator colour was dark-blue at indicator onset (Go 100%, Stop 0%), participants immediately knew that they could focus on the Go task (anticipate and respond at the target line, 800 ms from onset). Lighter blue colours signalled an increasing likelihood that a trial would be a Stop trial i.e. medium-blue (Go 75%, Stop 25%) and light-blue (Go 66.66%, Stop 33.33%). To measure reactive inhibition, two separate staircasing algorithms ensured convergence towards 50% success on Stop trials for the medium-blue and light-blue conditions.

Participants first completed two blocks of practice trials. The first block consisted of 60 Go trials, equiprobable across the three conditions (dark-blue, medium-blue, light-blue). The participant was instructed to focus solely on accurate Go task performance and that the colour of the bar was not informative for this block. The second practice block again consisted of 60 trials, equiprobable across the three conditions. Go and Stop trials occurred according to the frequencies outlined above. Participants were instructed that now the indicator might stop automatically prior to the target, and that when this happened, their goal was to inhibit their response i.e. prevent releasing the switch (Stop trials). Participants were informed that the majority of trials would be Go trials, that Stopping would not always be possible due to a stop-signal delay tracking algorithm, and that they should perform to their best ability on both Go and Stop trials. To reinforce the Go task requirements, the colour of the target line changed to green, yellow, orange or red at the end of each trial, depending on whether the response was within 20, 40, 60, or >60 ms of the target.

Then, the two main EEG blocks were completed (260 trials per block, 520 trials in total). Each block comprised 60 trials for which the indicator colour was dark-blue (Go_100_), 80 trials for which the indicator colour was medium-blue (Go_75_ | Stop_25_), and 120 trials for which the indicator colour was light-blue (Go_66_ | Stop_33_). Response time was recorded relative to the target on each trial, and the indicator was reset to empty after 1000 ms. The inter-trial interval to the next warning cue was 750 ms.

The two task runs were merged into a single behavioural data set per session. Race model assumptions were checked to ensure that p(respond|signal) was between 0.25 and 0.75 for all participants, sessions, and conditions. For each participant and condition, response time on p(respond|signal) trials, i.e. Stop Fail trials, was also compared to Go trials to check the independence assumption. The data for one participant violated this race model assumption and therefore this participant was omitted from behavioural and EEG analyses of Stop trial data. SSRT was calculated using the integration method, with Go omissions replaced by maximum RT (1000ms) [Verbruggen et al., 2019].

### EEG acquisition and pre-processing

Continuous EEG was acquired from 60 scalp electrodes using a BrainAmp DC system (Brain Products) digitized at 2500 Hz. Electrode locations were per the international 10-20 system via a standard 64 Channel actiCAP (Brain Products), with the ground electrode at AFz and the reference at FCz for data recording. Channels FT9, FT10, TP9, and TP10 were omitted from EEG recording and reallocated to bipolar vertical and horizontal electrooculogram recordings (VEOG and HEOG).

Data were processed using Matlab r2017a (MathWorks) scripts implementing EEGLAB v14.1.1 (Delorme and Makeig, 2004) and ERPLAB v7.0.0 [Lopez-Calderon, 2014] routines. Data pre-processing involved merging the two runs into a single data file, automatic detection of bad channels using kurtosis (threshold 5) followed by visual inspection, spherical interpolation of bad channels, extraction of epochs, baseline correction using the 500 ms prior to each event, re-referencing to the average of all channels (excluding the VEOG and HEOG channels), and two-rounds of semi-automated artefact rejection. The first round identified and rejected trials with large muscle artifacts using a moving window applied to all EEG channels (threshold 200 uV; window width = 200 ms; window step = 50 ms). The second round identified and excluded trials containing blinks in the bipolar VEOG channel (threshold = 50 uV; window width = 400 ms; window step = 10 ms). Details of trial rejection, including the percentage of trials excluded, are provided in the Supplementary Materials (Tables 1 and 2).

**Table 1.**
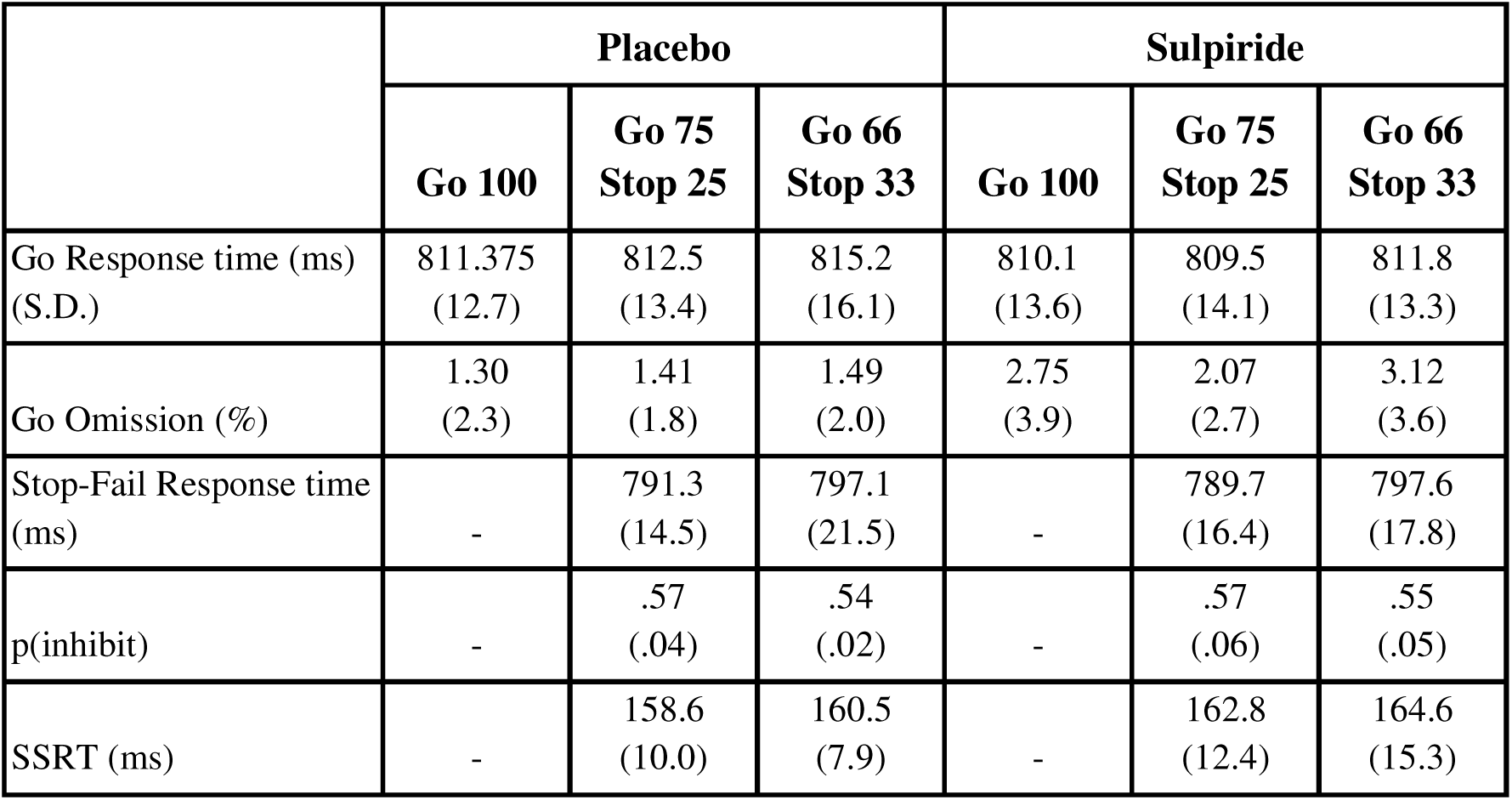
Behaviour. Mean ± Standard Deviation.

### Analyses – Event-related potentials

The ERP analyses comprised: i) An *onset-locked analysis* i.e. time-locked to the onset of the indicator fill (−500ms to +1000ms), with epochs baseline corrected using the 500 ms window interval preceding the stimulus onset. ii) A *stop-locked analysis* i.e. time-locked to the onset of the Stop cue (−500ms to +500ms). For this stop-locked analysis, a virtual Stop event was created for each Go trial as follows: For the dark-blue (Go_100_) condition, virtual event markers were generated by subtracting the participant’s SSRT (averaged across the two Stop conditions) from their Go response time on each individual trial. For the medium-blue (Go_75_) and light-blue (Go_66_) conditions, virtual event markers were generated by subtracting the participant’s SSRT per condition from Go response time on each individual trial. The analyses resulted in 5 trial-types: Go_100_; Go_75_; Go_66_; Stop_25_; Stop_33_. For the stop-locked analysis, baseline correction was over the 500 ms prior to the Stop-signal onset (virtual Stop-signal onset for Go trials). For both onset-locked and stop-locked analyses, averaged ERPs were filtered with a 6th order bandpass Butterworth filter between 0.1Hz and 40Hz.

Statistical analyses were performed using the MATLAB toolbox BRAINSTORM (version 3.26). For ERP analyses, two-tailed cluster-based permutation tests were conducted using the *ft_timelockstatistics* function from the FieldTrip plugin. The minimum number of neighbouring channels per cluster was set to two, with significance threshold (α) set at 0.05. For the onset-locked analysis assessing proactive inhibition, statistical testing was performed over the 0-800 ms interval following stimulus onset to compare ERP’s for: i) Go Probability (Go_100_, Go_75_, Go_66_) within each session, and ii) Go across Drug (Placebo vs Sulpiride). For the stop-locked analysis, assessing reactive inhibition, the test was conducted over the 0-500 ms interval following the stop-signal onset (Go vs Stop Success).

Finally, for the stop-locked data, difference ERPs were also calculated for each participant by subtracting the successful Stop ERPs from Go ERPs (for the corresponding condition). The Go ERP was subtracted to better isolate the ERP waveform related to the Stop process from the ongoing movement preparation process. These difference ERPs were then used for permutation t-tests (settings as described above) to assess for any effect of Drug (Placebo vs Sulpiride) on reactive inhibition to the stop cue.

### Analyses - Behaviour

Behavioural data were analysed with repeated measures analysis of variance (rmANOVA). To assess proactive inhibition, mean Go response time was analysed with 2×3 rmANOVA, factors Drug (Placebo, Sulpiride) and Go Probability (Go_100_, Go_75_, Go_66_). To assess variability in timing of the anticipated response, Go response time variability (standard deviation of the Go response distribution) was similarly analysed with 2×3 rmANOVA. We also verified, at the group level, that response times on Stop-Fail trials were faster then Go trials with a 2×2×2 rmANOVA, factors Drug, Go Probability (Go_75_, Go_66_), and Condition (Go, Stop-Fail). To assess reactive inhibition, SSRT was interrogated with 2×2 rmANOVA, factors Drug, and Condition (Stop_25_, Stop_33_).

JASP v0.12 was used for statistical analysis of behaviour. For parametric statistics, α was set at .05, and effect size is reported as Cohen’s d or partial. Where appropriate, main effects and interactions were followed by Bonferroni-corrected post-hoc tests. Analogous Bayesian rmANOVAs were also conducted to determine evidence in favour of the alternative hypothesis (BF_10_) with effects reported across matched models (BF_incl_), and BF_incl_ values >1 interpreted as weak evidence, >3 moderate evidence, and >10 strong evidence.

## Results

Twenty-four (N=24) healthy volunteers with normal or corrected-to-normal vision were included in this study. The sample was aged 18-39 years (*M* = 24.1 years, *SD* = 6.3), predominantly male (91%), and right handed according to the Edinburgh Handedness Inventory (*M* = +91.9, *SD* = 6.2). Importantly, the session in which sulpiride was administered was correctly guessed by 54% of participants, which was no better than chance (Χ^2^ = 0.17, *p* = .68), indicating successful blinding to drug condition. The mean confidence rating of participants who correctly guessed the sulpiride session was 62% (*SD* = 19%) and the mean confidence rating of participants who did not guess correctly was 66% (*SD* = 20%), indicating no systematic difference in confidence ratings (*p* = .98).

### Behaviour

Figure 2 summarises performance on the task and Table 1 shows mean and standard deviation of Go response times. There was no main effect of Drug on mean Go response time (*F*_1,22_ = 1.45, *p* = .24, *η*^2^*_p_* = .06, BF_incl_ = 1.44) or a Go probability by Drug interaction (*F*_2,44_ = 0.97, *p* = .39, *η*^2^*_p_* = .04, BF_incl_ = 0.15). There was, however, a main effect of Go Probability (*F*_2,44_ = 3.66, *p* = .03, *η*^2^*_p_* = .14, BF_incl_ = 0.44) with mean Go response time increasing as the likelihood of a Stop cue increased (Figure 2A). Simple main effects analysis indicated a significant main effect of Go Probability for Placebo (*F*_2_ = 4.15, *p* = .02), consistent with previous studies of proactive inhibition [Leunissen et al., 2016; Zandbelt and Vink, 2010], but not Sulpiride (*F*_2_ = 1.41, *p* = .25).

**Figure 2.**
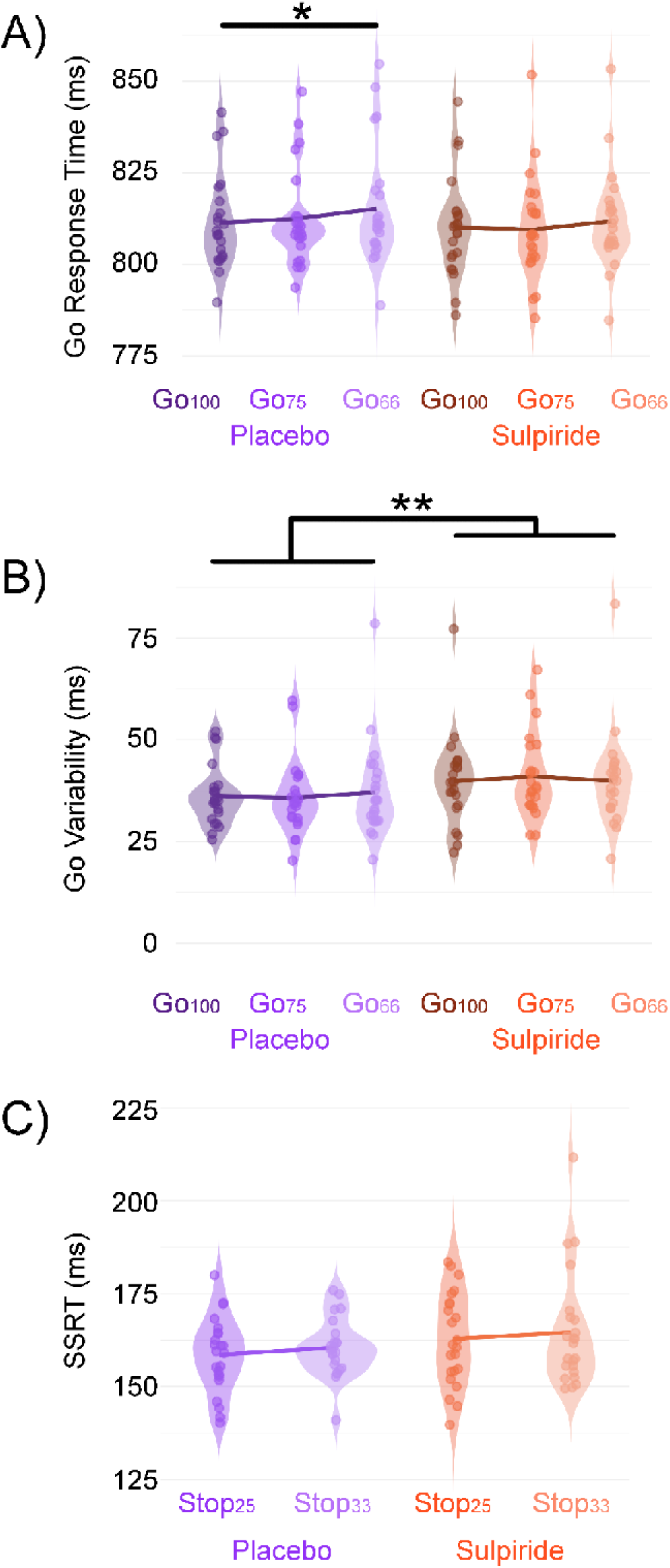
Behavioural results. (A) Main effect of Go condition on Go response time. Simple main effects analysis was significant for Placebo but not Sulpiride. (B) Main effect of Drug session on Go response time variability (Go variability determined as the standard deviation of the Go response time distribution for each participant). (C) Stop-signal reaction time (SSRT) for Placebo and Sulpiride sessions.

Drug increased the variability of the Go response time distribution (*F*_1,22_ = 7.79, *p* = .01, *η*^2^*_p_* = .26, BF_incl_ = 128.11) with the bayes factor indicating strong evidence of greater response variability for Sulpiride compared with Placebo (Figure 2B, Table 1). There was no main effect of Go Probability (*F*_2,44_ = 0.10, *p* = .90, *η*^2^*_p_* = .005, BF_incl_ = 0.07), or Drug x Go Probability interaction (*F*_2,44_ = 0.64, *p* = .53, *η*^2^*_p_* = .03, BF_incl_ = 0.19).

In alignment with race model assumptions, response time was significantly faster on failed Stop trials, i.e. p(respond|signal), compared with Go trials (Table 1). There was a main effect of Condition (*F*_1,22_ = 153.97, *p* < .001, *η*^2^*_p_* = .88, BF_incl_ = 116.7 × 10^24^, extreme evidence) with responses occurring earlier on StopFail trials compared to Go trials, a main effect of Go Probability (*F*_1,22_ = 11.49, *p* = .003, *η*^2^*_p_* = .34, BF_incl_ = 44.57, strong evidence) with later responses for the Go_66_ condition compared to the Go_75_ condition, and a Condition x Go Probability interaction (*F*_1,22_ = 4.82, *p* = .04, *η*^2^*_p_* = .18, BF_incl_ = 0.73). The main effect of Drug and interactions involving Drug were not significant (*all F* < 2.56, *p* > .12, BF_incl_ < 0.42).

Stop signal reaction time, SSRT, showed a 4 ms increase for Sulpiride compared with Placebo (Figure 2D, Table 1), with no significant main effects or interactions (Drug main effect *F*_1,22_ = 3.91, *p* = .06, *η*^2^*_p_* = .15, BF_incl_ = 1.62, weak evidence; Go Probability *F*_1,22_ = 0.91, *p* = .35, *η*^2^*_p_* = .04, BF_incl_ = 0.33; Drug x Go Probability *F*_1,22_ = 0.002, *p* = .91, *η*^2^*_p_* = .00, BF_incl_ = 0.28).

### Go trial event-related potentials (ERPs) time locked to indicator onset

The onset-locked analysis focussed on a slowly developing negativity centred on frontocentral electrodes during Go trials. The grand average topographies for each session are shown for each Go condition (Figure 3A).

**Figure 3.**
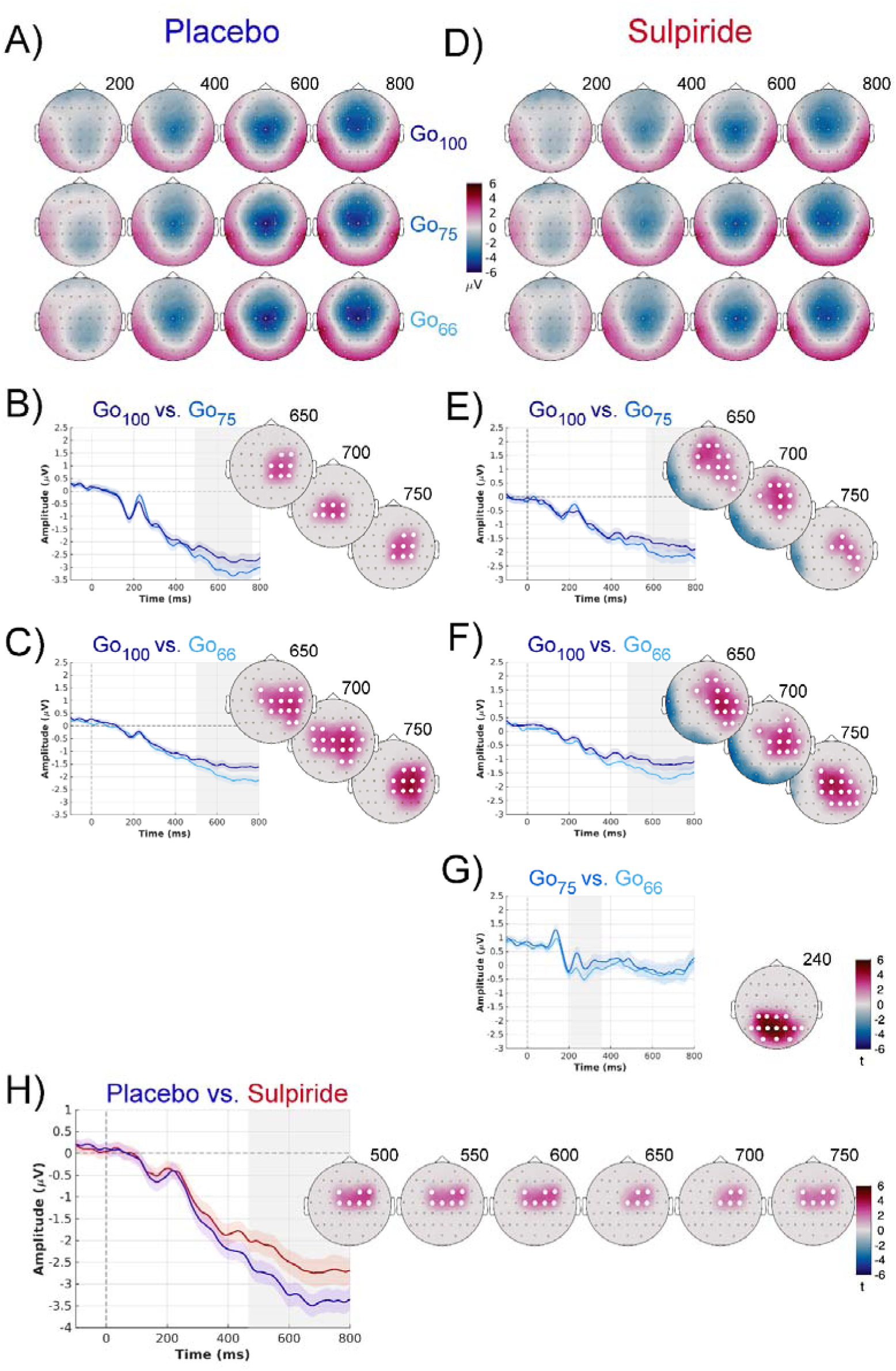
Go-trial onset-locked ERP analysis. (A-C) Scalp topographies of the Go-locked ERP for the GO_100_, GO_75_, and GO_66_ conditions at representative time points in the Placebo session. ERP waveforms and corresponding statistical topographies are shown for the contrasts GO_100_ vs. GO_75_ (p = 0.01) and GO_100_ vs. GO_66_ (p = 0.006), respectively. Statistical topographies display t-values, with the significant electrode cluster highlighted. ERP waveforms were obtained by averaging activity across electrodes within the significant cluster. Shaded regions indicate the time windows identified as significant by the cluster-based permutation test. (D-G) Scalp topographies of the Go-locked ERP for the GO_100_, GO_75_, and GO_66_ conditions in the Sulpiride session at representative time points. ERP waveforms and corresponding statistical topographies for the contrasts GO100 vs. GO75 (p = 0.01), GO100 vs. GO_66_ (p = 0.006), and GO_75_ vs. GO_66_ (p = 0.006), respectively. Statistical topographies represent t-values, with significant electrode clusters highlighted. ERP waveforms were generated by averaging activity across electrodes within the corresponding cluster, and shaded regions denote significant temporal clusters. (H) ERP waveform and statistical topography for the Placebo vs. Sulpiride contrast examining the main effect of the Drug, with all Go conditions collapsed. A cluster-based permutation test identified a significant positive centrofrontal cluster (p = 0.03, uncorrected).

Within the Placebo session, cluster-based permutation tests revealed significant positive clusters for the Go_100_ versus Go_75_ contrast (p = 0.010, 487–763 ms; Bonferroni-corrected) and the Go_100_ versus Go_66_ contrast (p = 0.006, 499–800 ms; Bonferroni-corrected) (Figure 3A-C).

Within the Sulpiride condition, significant positive clusters were observed over a similar time window for both the Go_100_ versus Go_75_ contrast (p = 0.006, 564–772 ms; Bonferroni-corrected) and the Go_100_ versus Go_66_ contrast (p = 0.002, 478–800 ms; Bonferroni-corrected) (Figure 3D-F). In addition, the Go_75_ versus Go_66_ contrast revealed a significant positive cluster with a posterior topography in an earlier time window (p = 0.006, 206–360 ms; Bonferroni-corrected) (Figure 3G).

Finally, to examine the main effect of Drug, all Go conditions were collapsed, and the Placebo vs. Drug contrast was assessed. This analysis revealed a significant cluster around 432–800 ms (p = 0.03, uncorrected; Figure 3H).

### Event-related potentials (ERPs) time locked to the stop-cue

For the stop-locked ERP analysis, significant clusters were identified in both the Placebo and Sulpiride conditions. For the Stop_25_ versus Go_75_ contrast, a significant positive cluster was observed in the Placebo condition from 79–346 ms (p = 0.010; Bonferroni-corrected), and in the Sulpiride condition from 0–349 ms (p = 0.006; Bonferroni-corrected) (Figure 4 A-B). The significant time window included a prominent P1 potential for Stop that preceded the SSRT, along with subsequent N2 and P3 peaks that followed the SSRT.

**Figure 4.**
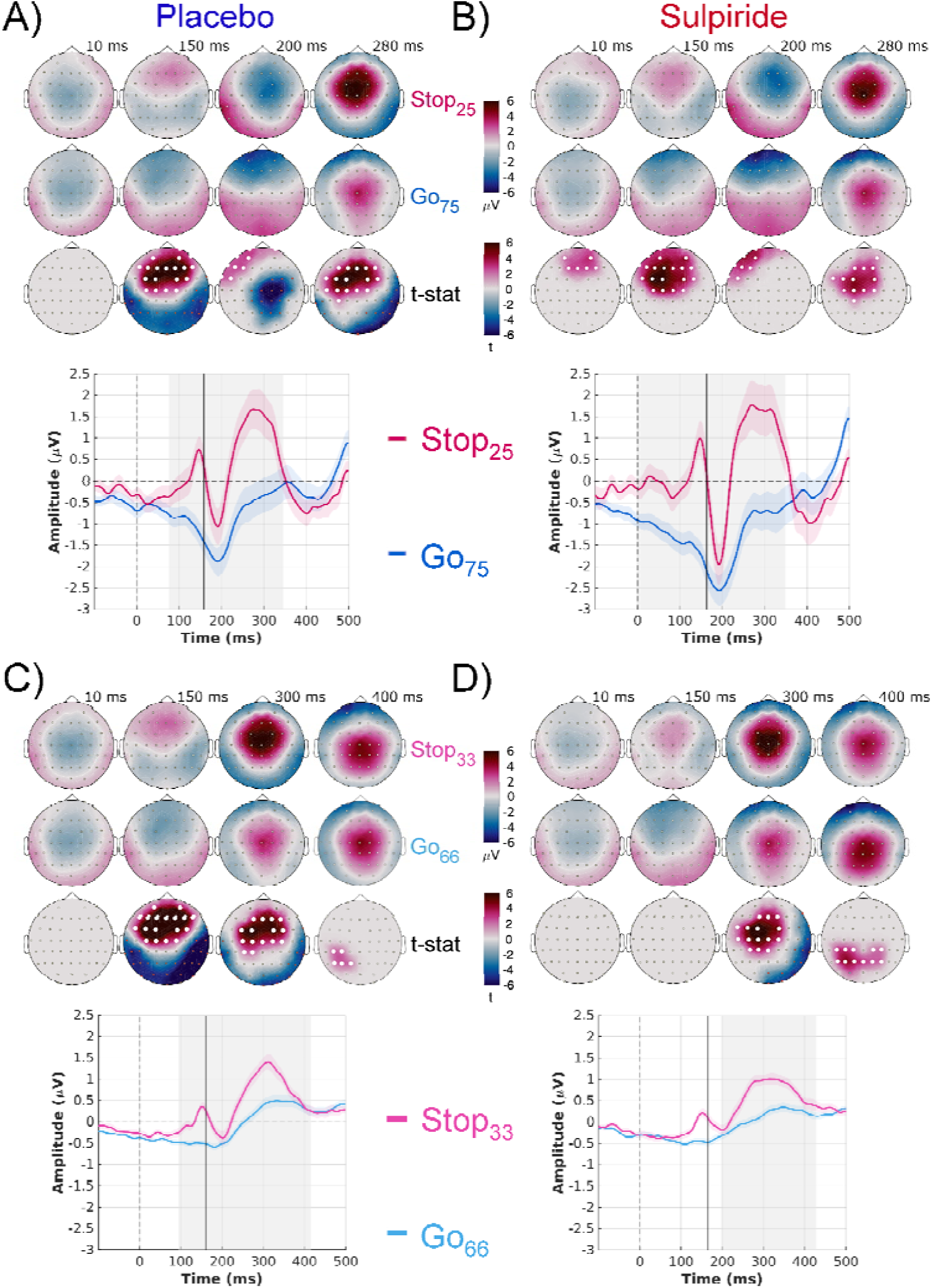
Stop-locked ERP analysis for Go and Stop conditions. (A) Placebo ERP waveforms and scalp topographies for the STOP_25_ vs. GO_75_ contrast (p = 0.01, Bonferroni-corrected). ERP waveforms were generated by averaging activity across electrodes within the significant positive frontocentral cluster. The first two rows of topoplots show scalp activity at representative time points, while the third row displays t-statistic maps for the contrast, with electrodes belonging to the significant cluster highlighted. (B) Sulpiride ERP waveforms and statistical topographies for the STOP_25_ vs. GO_75_ contrast (p = 0.006, Bonferroni-corrected), revealing a stronger positive frontocentral cluster than in the Placebo condition. (C) Placebo ERP waveforms and statistical topographies for the STOP_33_ vs. GO_66_ contrast (p = 0.008), with the significant frontocentral cluster highlighted. (D) Sulpiride ERP waveforms and statistical topographies for the STOP_33_ vs. GO_66_ contrast (p = 0.004), showing a significant positive frontocentral cluster. In all panels, shaded regions indicate the significant temporal clusters identified by the cluster-based permutation test. The solid vertical line marks the group-mean SSRT. Note the prominent P1 peak that precedes the SSRT.

For the Stop_33_ versus Go_66_ contrast, both groups showed significant positive clusters, with the Placebo condition exhibiting a cluster spanning 95–415 ms (p = 0.008; Bonferroni-corrected) and the Sulpiride condition exhibiting a cluster spanning 196–428 ms (p = 0.004; Bonferroni-corrected) (Figure 4 C-D). Again, there was a prominent P1 potential for Stop that preceded the SSRT, along with a subsequent N2 and P3 peak that followed the SSRT.

To examine the main effect of Drug on reactive inhibition, independent of activity related to movement preparation, we first computed the difference ERP between Stop and Go trials (Stop − Go) separately for the Placebo and Sulpiride sessions. We then contrasted these difference waveforms between the two drug conditions using a cluster-based permutation test. No significant clusters were identified for either the Stop_25_ vs. Go_75_ or Stop_33_ vs. Go_66_ comparisons, indicating that sulpiride did not significantly modulate the stop-locked ERP relative to Placebo (Figure 5).

**Figure 5.**
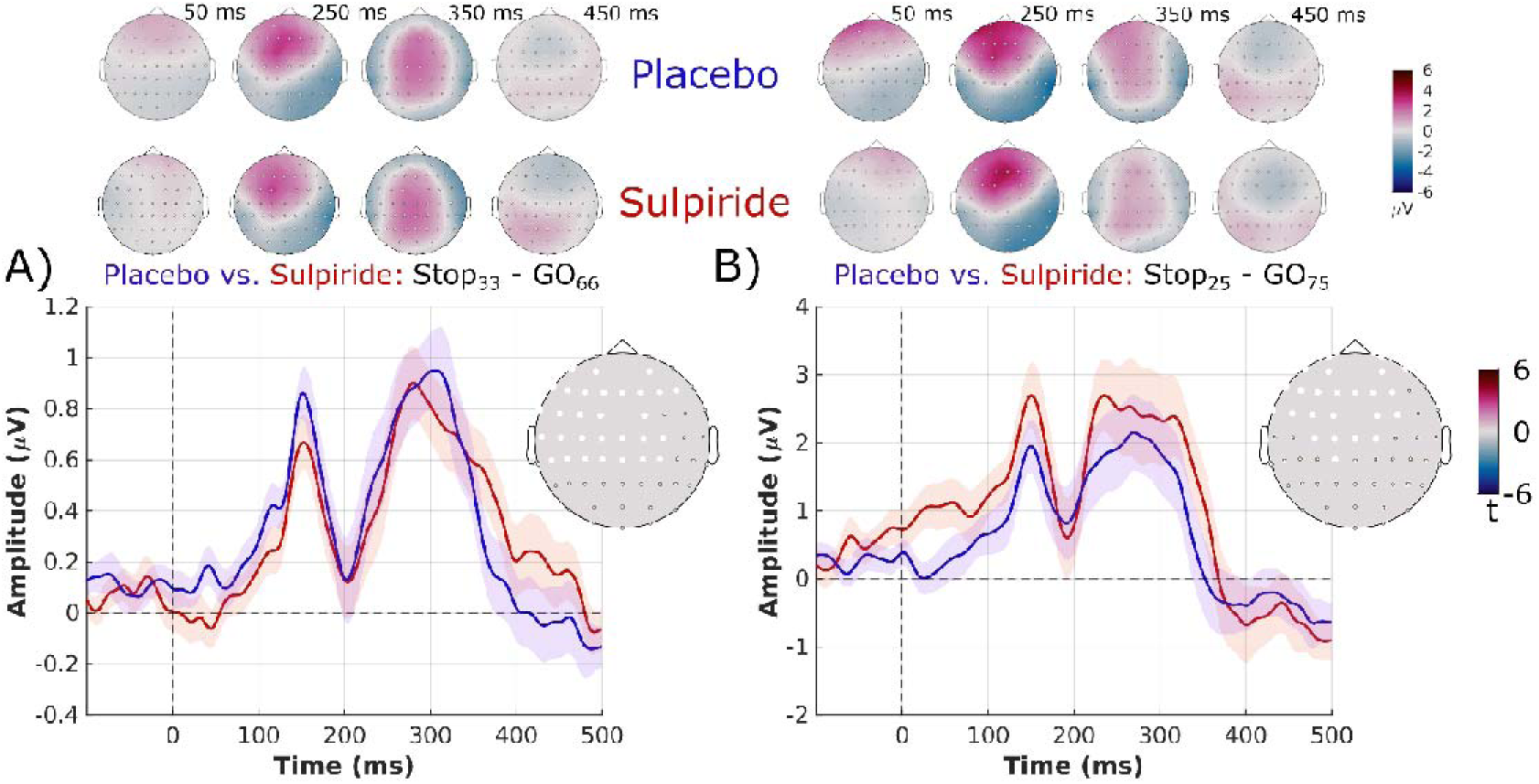
Comparison of stop-related difference ERP waveforms reveals no main effect of Drug on reactive inhibition. (A) STOP_33_ - GO_66_ difference ERP waveforms for the Placebo and Sulpiride conditions. The top two rows show scalp topographies of the difference waveforms (STOP_33_ - GO_66_) at representative time points for each drug condition. (B) STOP_25_ - GO_75_ difference ERP waveforms for the Placebo and Sulpiride conditions. The top row shows scalp topographies of the difference waveforms (STOP_25_ - GO_75_) at representative time points for each drug condition. ERP waveforms for both plots were generated by averaging activity across common frontocentral electrode clusters defined as the overlap of the significant clusters identified in the STOP_25_ vs. GO_75_ and STOP_33_ vs. GO_66_ analyses within the Placebo and Sulpiride conditions. The t-stat topoplot represents the averaged electrodes for each ERP. No significant differences were observed between the Placebo and Sulpiride conditions in either comparison, as revealed by the cluster-based permutation test results.

## Discussion

The present results indicate that sulpiride did not have a strong impact on proactive inhibition at the level of mean Go response times. Increasing the likelihood of Stop trials had the expected effect of increasing Go response times in the placebo session but not the sulpiride session, in the absence of a significant interaction. However, sulpiride resulted in increased variability of Go response times and an attenuated amplitude of the frontocentral negativity during movement preparation, compared with placebo. Additionally, although there was a trend toward longer SSRTs with sulpiride (∼4 ms), this difference was not statistically significant. Sulpiride also did not modulate stop-locked reactive inhibition ERPs compared with placebo. These results indicate that blocking of dopamine D2 receptors by sulpiride administration influenced processes associated with proactive inhibition, potentially via an effect on the indirect basal ganglia pathway, but left reactive inhibition unchanged.

### Effects of D2 blockade in relation to the hyperdirect, direct, and indirect basal ganglia pathways

A non-significant change in SSRT is contrary to our hypothesis that sulpiride would impair reactive inhibition based on work in rodents reporting that SSRT increased after sulpiride administration [Eagle et al., 2011]. On the one hand, studies show that indirect catecholamine agonists, such as methylphenidate, and D2 agonists, such as cabergoline improve reactive inhibition [Nandam et al., 2011a; Nandam et al., 2013]. These effects may depend on drug dose and baseline neurochemical state, with Koyun et al. [2025] showing that low doses of methylphenidate result in better inhibitory control, with higher baseline striatal gamma-aminobutyric acid (GABA) to glutamate ratios predicting better response inhibition accuracy. On the other hand, our findings and those of Rawji et al. [2020], indicate that pharmacological manipulation of D2/D3 receptors does not necessarily alter reactive inhibition. Thus the literature is mixed, and the evidence for D2 receptor involvement in reactive inhibition is equivocal in humans.

Reactive stopping is thought to be achieved via the hyperdirect pathway. The hyperdirect pathway bypasses the striatum, where the DRD2 receptors are highly expressed, projecting directly from the cortex to the STN [Nambu et al., 2002]. The non-significant effect of the drug on SSRT is consistent with the interpretation that reactive inhibition is implemented via the hyperdirect, as opposed to the indirect pathway, and may reflect species-related differences in task design and implementation. The indirect pathway has been suggested to play a greater role in proactive inhibition [Aron, 2011], supported by studies highlighting greater fMRI activation of the caudate head in trials with high stop-signal probability compared to low stop-signal probability [Leunissen et al., 2016]. Our finding of sulpiride altering proactive inhibition, by rendering Go response times less precise, further supports the involvement of the indirect pathway in proactive inhibition.

The direct and indirect pathways were thought to function independently of one another, with activation of the direct pathway involved in facilitation of actions and the indirect pathway in suppression of actions [Albin et al., 1989; DeLong, 1990]. However, studies have shown simultaneous activation of both direct and indirect pathways preceding movement initiation [Cui et al., 2013], challenging this notion. Instead, there may be dynamic competition between the indirect and direct pathways, with the relative activation of each shifting the overall network toward a facilitating or suppressive state [Dunovan and Verstynen, 2016]. This dynamic competition likely interacts with the hyperdirect pathway, which prevents execution of the selected action if recruited early enough [Dunovan and Verstynen, 2016]. Moreover, the competition between the direct and indirect pathway is suggested to be modulated by dopamine with a dependent process computational model showing that increasing dopamine levels result in quicker decisions and reduced inter-trial variability in reaction times, whereas lower levels of dopamine lead to high inter-trial variability in reaction times [Dunovan and Verstynen, 2016]. Our data is in line with this computational modelling work, wherein blockade of dopamine D2 receptors resulted in increased variability of Go response times. Essentially, the D1 and D2 dynamic competition that is responsible for setting the response threshold seems to be broken down by sulpiride. This also corroborates the findings of Rawji et al. [2020], wherein drift diffusion modelling highlighted that ropinirole, a D2/D3 receptor agonist, diminished the ability to raise the decision threshold under conditions requiring proactive inhibition. Dopamine thus seems to play an important role in influencing the extent to which proactive control is recruited.

### Proactive inhibition and frontocentral negativity

The CNV has been proposed as a marker of proactive inhibition, with stronger late CNV observed in high stop-signal probability conditions as compared to low stop-signal probability conditions [Brevers et al., 2020]. The results of the current study are in line with this. Although we did not find a significant difference in fronto-central negativity between Go_75_ and Go_66_, we found significant differences between Go_100_ and Go_75_ and between Go_100_ and Go_66_, with lower Go probability (i.e., higher stop-signal probability) trials showing greater negativity. The late CNV reflects processes related to motor preparation [Verleger et al., 2000]. A stronger CNV when proactive inhibition is greater, as reflected by increased mean Go response times, indicates that increased motor preparation is required to counteract proactive inhibition in high-stop signal probability conditions. Sulpiride made the Go response times more variable and also attenuated the negative potential, implying that it impaired some aspect of motor preparation. This is consistent with studies showing that depletion of striatal dopamine impairs response preparation in rats [Brown and Robbins, 1989].

### ERP components of Reactive Inhibition

We observed prominent N2 and P3 peaks following the stop-signal, replicating previous findings [Kok et al., 2004; Zhu et al., 2024]. The P3 component has been proposed to reflect an inhibitory control process and is considered essential for action-stopping [Hervault et al., 2025]. While some studies report that the P3 onset latency coincides with SSRT [Hervault et al., 2025], others show that the P3 onset latency is much longer than the SSRT [Huster et al., 2020; Thunberg et al., 2020]. In the current study, the P3 onset latency exceeded the estimated SSRT by >100 ms, for both the placebo and sulpiride sessions. This raises the question of whether the P3 component is causally involved in implementing reactive inhibition.

Notably, a prominent P1 component that preceded the SSRT was evident in our data. Few studies of response inhibition have reported a P1 component, with Wessel and Aron [2015] observing a prominent P1 in only a few participants following a stop-signal. In the current study, we found a prominent P1 component at the group-level, for stop-signal trials that preceded the estimated SSRT and the N2/P3 complex. The strong consistent P1 effect in the current study is likely an advantage of the specific paradigm and analysis approaches, including the creation of virtual stop events for Go trials. The anticipated response paradigm in the current study provides a tighter control over the timing of the Go process (compared to reaction time versions of the stop-signal task), since the participant’s goal is to respond at a fixed point in time. Additionally, the task reinforces correct Go performance through trial-by-trial feedback on how close the indicator stopped relative to the target [Leunissen et al., 2017].

### Limitations

The current study addresses an important gap in the literature by examining the effects of sulpiride on proactive and reaction inhibition in humans, using a stop-signal task. Although our paradigm and analysis approach are key strengths of this study, a few limitations warrant consideration. The current study did not find an effect of sulpiride on mean Go response time (there was a simple main effect of Go response probability for placebo that was absent for sulpiride, however this did not manifest as a main effect or interaction). This is contrary to Rawji et al. [2020], who found that a D2/D3 agonist reduced proactive inhibition. One plausible explanation is that the sample size (N=23) provided insufficient statistical power to detect a small effect. Alternatively, the trial-by-trial feedback, conveyed through changes in colour of the target line to reinforce accurate timing of Go responses, may have encouraged participants to maintain consistent Go response times, on average, and thereby limited our ability to detect an effect of sulpiride on mean Go response time. Moreover, the relationship between dopamine and task performance is multifaceted, and often depends on an individual’s baseline levels of dopamine. Human PET studies have demonstrated an important role for basal dopamine in the striatum [Cools and D’Esposito, 2011]. Studies have also shown that basal dopamine neurotransmission interacts with drug effects. For example, ropinirole improved response inhibition in participants with low basal dopamine neurotransmission but impaired inhibition in adults with higher basal neurotransmission [MacDonald et al., 2016]. In a study by Eisenegger et al. [2014], sulpiride impaired choice performance and this effect was greater in subjects with a lower density of dopamine D2 receptors in the striatum. In the current study, we did not take genetic risk scores into account. Future studies should investigate how basal dopamine neurotransmission levels impact proactive inhibition after sulpiride administration. Finally, while sulpiride has a high affinity to D2 receptors, it also has some impact to block D3 and D4 receptors [Caley and Weber, 1995]. Therefore, we cannot conclude definitively that the effects of sulpiride on proactive response inhibition were mediated specifically by D2 receptors.

## Conclusion

To conclude, we have shown that sulpiride, a D2 antagonist, interferes with covert processes associated with response inhibition and motor preparation in humans. Sulpiride led to greater variability in Go response times and an attenuated frontocentral negativity, thought to be an electrophysiological marker of proactive inhibition. Sulpiride had no effect on reactive inhibition, reflected by no significant changes in SSRT, and did not alter ERP components associated with reactive stopping. Altogether, the pattern of results observed in this study are consistent with the hyperdirect pathway being critical for reactive inhibition and the indirect pathway for proactive inhibition.

## Statements and Declarations

### Ethical considerations

All procedures in the current study were approved by the Monash University Human Research Ethics Committee, Monash University, Melbourne (Ethics approval number: CF16/1572-2016000821). All ethical guidelines were in accordance with the Declaration of Helsinki.

### Consent to participate

Written informed consent was obtained from all participants.

### Data Availability Statement

The datasets generated during and/or analyzed during the current study are available from the corresponding author on request, subject to ethics approval requirements.

