## Supplementary Material for "Effect of dopamine D2 blockade on behavioural and electrophysiological measures of inhibitory control"

***Table 1: Percentage of trials excluded in EEG onset locked analysis***

| **Condition** | **Placebo % excluded (S.D.)** | **Sulpiride % excluded (S.D.)** |
| --- | --- | --- |
| Go100 | 20.1 (19.5) | 29.3 (23.1) |
| Go75 | 21.7 (21.5) | 31.4 (24.7) |
| Go66 | 21.2 (21.4) | 29.8 (24.2) |
| Stop25 | 17.2 (19.3) | 29.5 (24.2) |
| Stop66 | 21.8 (28.1) | 30.9 (24.6) |

***Table 2: Percentage of trials excluded in EEG stop locked analysis***

| **Condition** | **Placebo % excluded (S.D.)** | **Sulpiride % excluded (S.D.)** |
| --- | --- | --- |
| Go100 | 11.8 (15.7) | 21.3 (19.2) |
| Go75 | 11.3 (15.7) | 21.7 (20.6) |
| Go66 | 11.5 (15.6) | 20.7 (19.9) |
| Stop25 | 10.8 (15.3) | 21.1 (18.5) |
| Stop66 | 12.5 (20.0) | 24.1 (20.4) |

For Table 1 and 2, the percent trials excluded are reported following the two rounds of artifact rejection described in the methods.

The main analyses reported in the manuscript make use of contemporary cluster based permutation testing to identify the significant electrodes and time windows. To aid comparison with older literature reporting ERPs for stop-signal tasks, here we provide figures for the data averaged over electrodes FCz and Cz, which are in close alignment with the figures presented in the main manuscript.

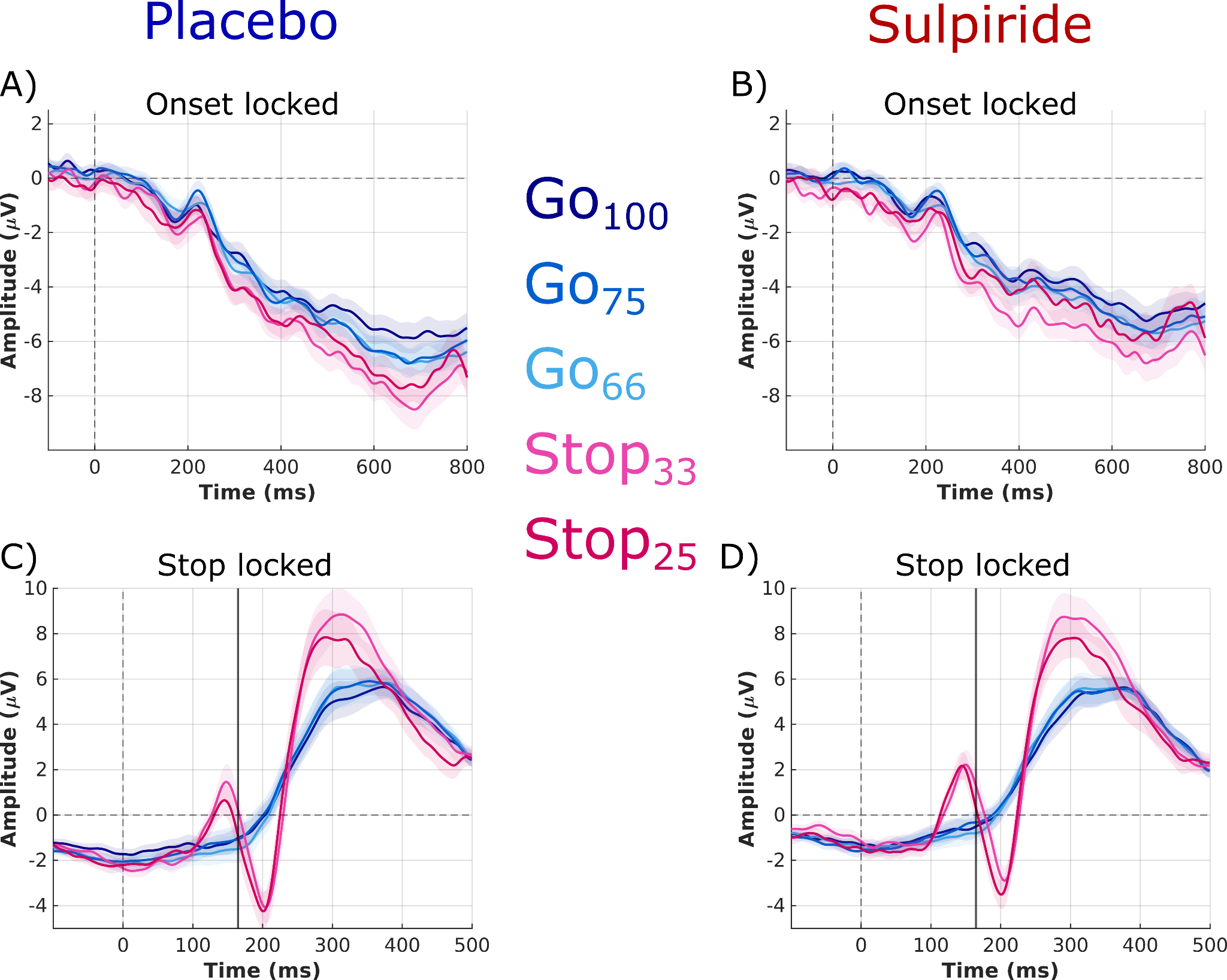

***Supplementary Figure 1.* Onset-locked and stop-locked ERP waveforms for Go and Stop conditions averaged across electrodes FCz and Cz.** (A) Placebo ERP waveforms time-locked to the onset of indicator fill for the GO_100_, GO_75_, and GO_66_, STOP_33_ and STOP_66_ conditions. (B) Sulpiride onset-locked ERP waveforms for the GO_100_, GO_75_, and GO_66_, STOP_33_ and STOP_66_ conditions. (C) Placebo ERP waveforms time-locked to the onset of the stop-cue for STOP_33_ and STOP_66_ conditions and to the virtual stop-cue for GO_100_, GO_75_, and GO_66_, STOP_33_ and STOP_66_ conditions. (D) Sulpiride stop-locked ERP waveforms for the GO_100_, GO_75_, and GO_66_, STOP_33_ and STOP_66_ conditions. In the stop-locked analysis panels, the solid vertical line marks the group-mean SSRT. Note the prominent P1 peak that precedes the SSRT and subsequent N2 and P3 peaks.
